## Supplementary material for "Reference Sequence Browser: An R application with a user-friendly GUI to rapidly query sequence databases": S1 Protocol

RESERVED DOI:
**10.17504/protocols.io.q26g71zxqgwz/v1**

Sriram Ramesh[1], Samuel Rapp[1], Jorge Tapias Gomez[2], Zia Truong[1], Dickson Chung[1], Benjamin Levine[1], Daniel Tapias-Gomez[3]

[1]University of California Santa Cruz; [2]Cornell University; [3]University of Texas Southwestern

Jorge Tapias Gomez
Cornell University

**Protocol Info:** Sriram Ramesh, Samuel Rapp, Jorge Tapias Gomez, Zia Truong, Dickson Chung, Benjamin Levine, Daniel Tapias-Gomez . Reference Sequence Browser: An R application with a User-Friendly GUI to rapidly query sequence databases. **protocols.io** **https://protocols.io/view/reference-sequence-browser-an-r-application-with-a-dazf2f3n**

**Created:** March 20, 2024

**Last Modified:** July 23, 2024

**Protocol Integer ID:** 97031

**Keywords:** environmental DNA, metabarcoding, shiny, NCBI, BOLD , CRUX, data access, reference sequence

### Abstract

Knowledge of which organisms have publicly available reference sequences at known DNA barcoding loci is crucial for both metabarcoding studies and the design of new primers. In a metabarcoding study, taxa of interest can not be detected if there are no labeled reference sequences to compare to. Similarly, designing species-specific primers is only possible if there are enough reference sequences for both the target taxa and their phylogenetic relatives to represent the genetic diversity across individuals. The purpose of this is to tool allow scientists to automatically gather reference sequences from NCBI, BOLD and CRUX. It is important to note that this is just a in-depth guide on how to use the tool, for more details on the features and their importance please refer to our paper in:
https://www.biorxiv.org/content/10.1101/2023.09.20.558722v3

### Guidelines

Please read this guide carefully to understand how to navigate and make the best use of the tool.

### Materials

There are no materials need other than a computer that can either run R locally and can download our tool or a stable internet connection to connect to our server.

### Safety warnings

We use the NCBI and BOLD APIs to query the respective databases and provide you with these results. We have seen these APIs stop working for brief periods of time, in particular BOLD. It isn't very common but it has happened before and could happen in the future.

In particular, the BOLD API presents certain constraints. Primarily, it enforces a rate limit based on the total number of species queries within an hour, capping the searches at around 100 species. Secondly, although rare, discrepancies have been observed where some organisms yielding incorrect metadata. In such instances, the affected organism are placed in the Absent/Invalid Metadata table described in **Step 16** and the keyword *invalid* appears alongside the organism. For accurate information, users are encouraged to manually verify these species directly through the database.

### Installation

1 The RSB is a Shiny GUI app shiny, which is available online at **https://sriramramesh.shinyapps.io/ReferenceSequenceBrowser/** or for download at https://github.com/SamuelLRapp/BlueWaltzBio. RSB uses the rentrez and bold CRAN packages to access the live NCBI and BOLD databases respectively and the Taxize package for some ease-of-use features.

Alternatively, the application can be installed locally. This has been tested for Windows, Mac-OSX, and Ubuntu. The tool uses R version 4.1.2. To install the app, download the zip file found here https://github.com/SamuelLRapp/BlueWaltzBio/releases/tag/v1.0.0-stable and then execute the script https://github.com/SamuelLRapp/BlueWaltzBio/blob/master/Coverage/rsbPackages.R to install the correct versions of the required libraries. The app can be run by opening an R console in the directory where the zip file was extracted and executing the command “shiny::runApp('Coverage')”.

| Software | |
|---|---|
| **Reference Sequence Browser** | NAME |
| Tested: Windows 10, MacOS Sonoma 14.1 | OS |
| BlueWaltzBio | DEVELOPER |
| Github | SOURCE LINK |

### Input File Structure

2 Throughout this guide you will see that all 3 pipelines allow you to use a CSV file to upload your species for the search. The format for uploading large inputs remains the same across all 3 databases.

There are two columns: “OrganismNames” and “Barcodes”. Every search tab uses the “OrganismNames” column, but “Barcodes” is only used by the NCBI Nucleotide search tab. Every pairing of entries in the “OrganismNames” and “Barcodes” columns will be searched for, so the two columns do not need to have the same number of elements.

We provide a CSV template for uploading data in our github in the https://github.com/SamuelLRapp/BlueWaltzBio/blob/master/Paper_Materials/paper_methods

_species.csv file that one can download to add their own species and barcodes.

### Welcome to the RSB

3 When you first start running the R Shiny application you will be presented with a welcome tab. In this tab, there are two main items that are important to pay attention to: the navigation bar and the NCBI input text box.

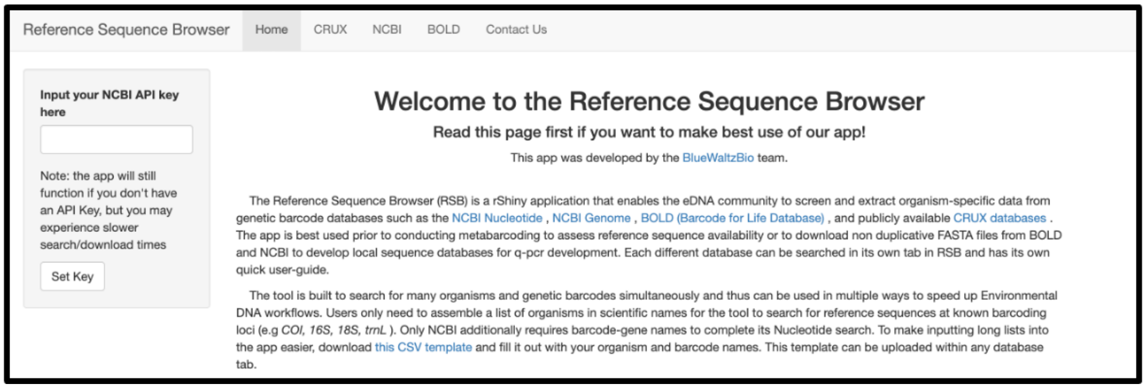


*Example of the RSB welcome tab that appears when you start running the R Shiny application*

4 The navigation bar at the top of the window is for navigating between databases. Currently, the tool supports 3 different databases: CRUX, NCBI Nucleotide, and BOLD.

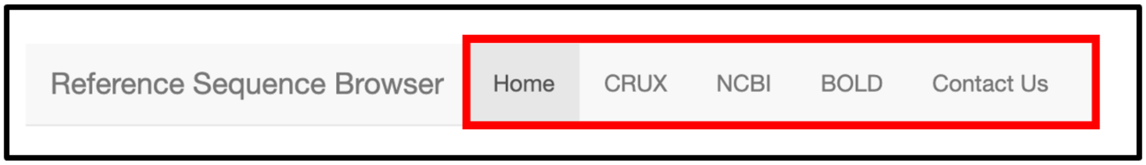


*Close up example of what the navigation bar looks like at the top of the window. The current tab is colored grey to indicate that is where we are currently located. In this case, that would be the Home Tab.*

5 Entering your own NCBI key into the API key input text box allows for quicker searches because the NCBI API is rate limited. Without a key, that rate is about 3 queries per second. With a key, that number goes up to 10 queries per second. For that reason, we highly recommend using an NCBI key. To create an NCBI key please go to: https://support.nlm.nih.gov/knowledgebase/article/KA-05317/en-us

5.1 To use an NCBI key, simply copy-paste the key into the text box titled "Input your NCBI API key here". Then click the "Set Key" button. Once this is done, there will be a pop up displaying whether the entered key was valid or not.

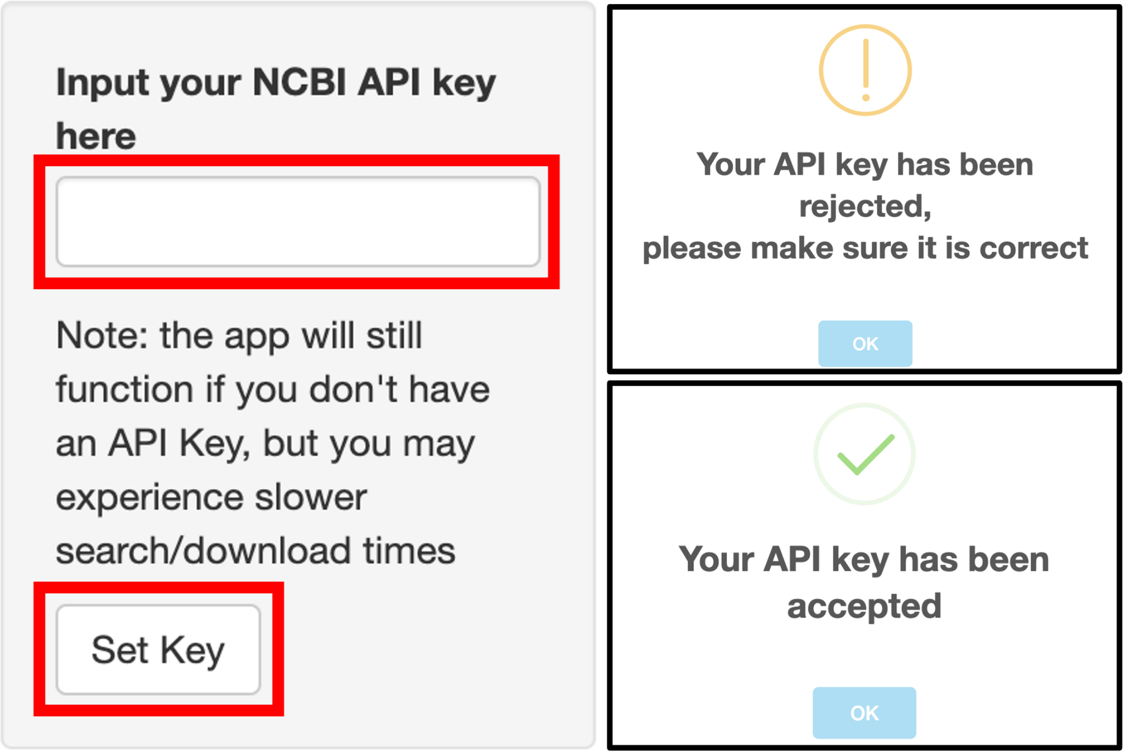


*Close up of the NCBI key input section, and the two possible pop-ups that appear depending on the whether the key was accepted or rejected.*

6 Now that we have covered the Welcome page of the RSB tool, we can move on to the databases.

### NCBI Tab

7 **Welcome Tab:** Using the navigation bar at the top, we can go to the NCBI section of the tool. Clicking on the NCBI button on the navigation bar will bring up the welcome tab of the NCBI database. There are several important things to pay attention to in this tab.

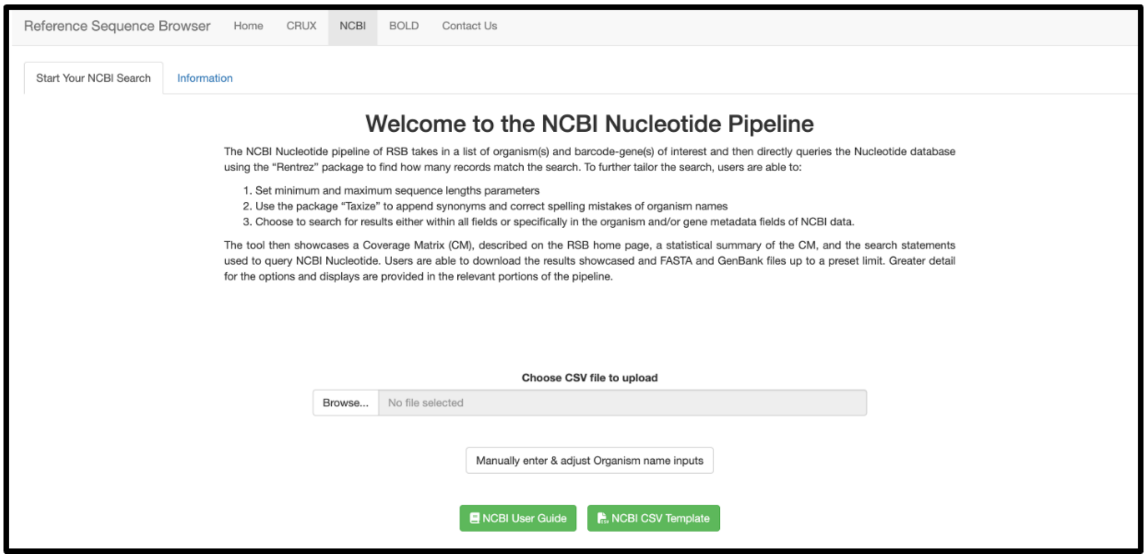


*Welcome section of the NCBI tab*

7.1 **Pipeline Navigation Bar:** The first thing to be aware of is that there is now a second navigation bar that will appear on the top left of the screen. This is a navigation bar for the database pipeline we have created and each tab is meant to be followed in a specific order (from left to right). When pulling up a pipeline for the first time in a session, only 2 tabs will be visible: the "Start your *Database* Tab" and an "Information Tab". Progressing through the pipeline will reveal more tabs, which can be clicked on to go back to previous steps.

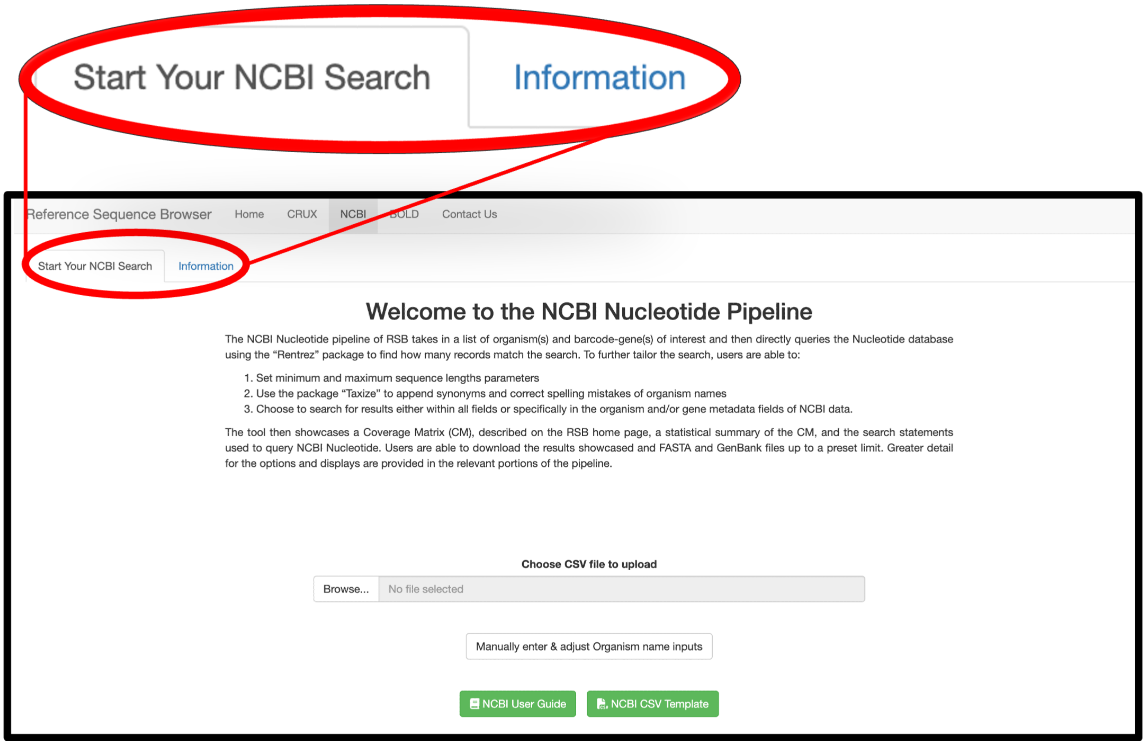


*Close up of the NCBI pipeline navigation bar*

7.2 **Buttons:** There are 4 main buttons in the NCBI welcome tab:

- **NCBI User Guide:** The NCBI User guide button links to this protocol.
- **NCBI CSV Template:** This button links to a google sheet. Download a copy of this sheet and fill it with any species and barcodes of interest. This CSV can be used to easily upload all the information to the tool instead of manually entering it later.
- **Browse:** Upload the filled CSV mentioned above by clicking on this button.
- **Manually enter & adjust Organism name inputs:** Click this button to move on to the next step.

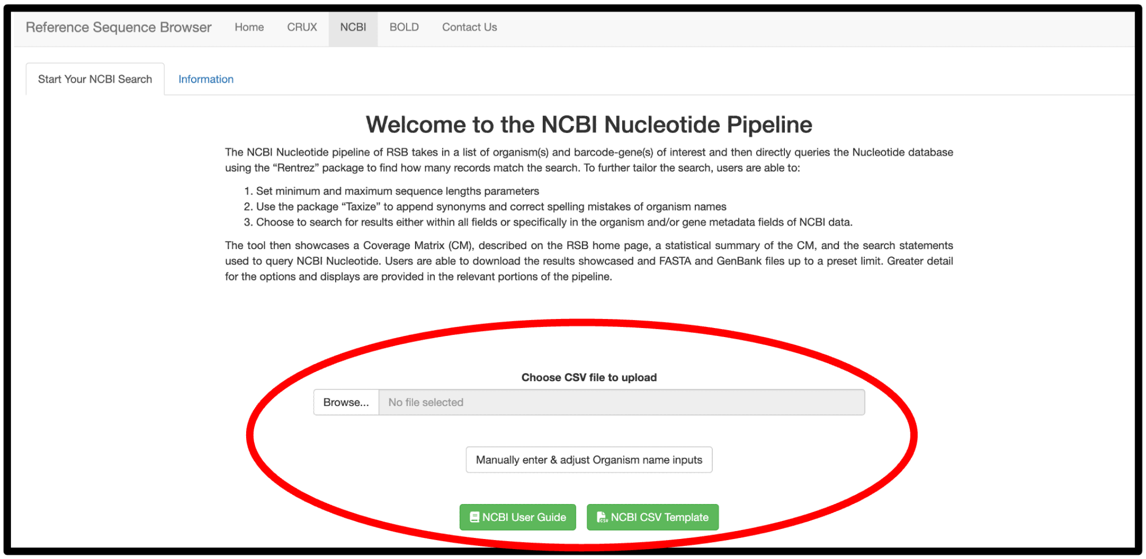


8 **Organism Names:** This screen is an opportunity to review and modify the organisms uploaded in the CSV, and toggle some search settings in relation to how the organisms will be searched for in the database. Once finished, click the "Manually enter & adjust Organism name inputs" button to proceed to the next step.

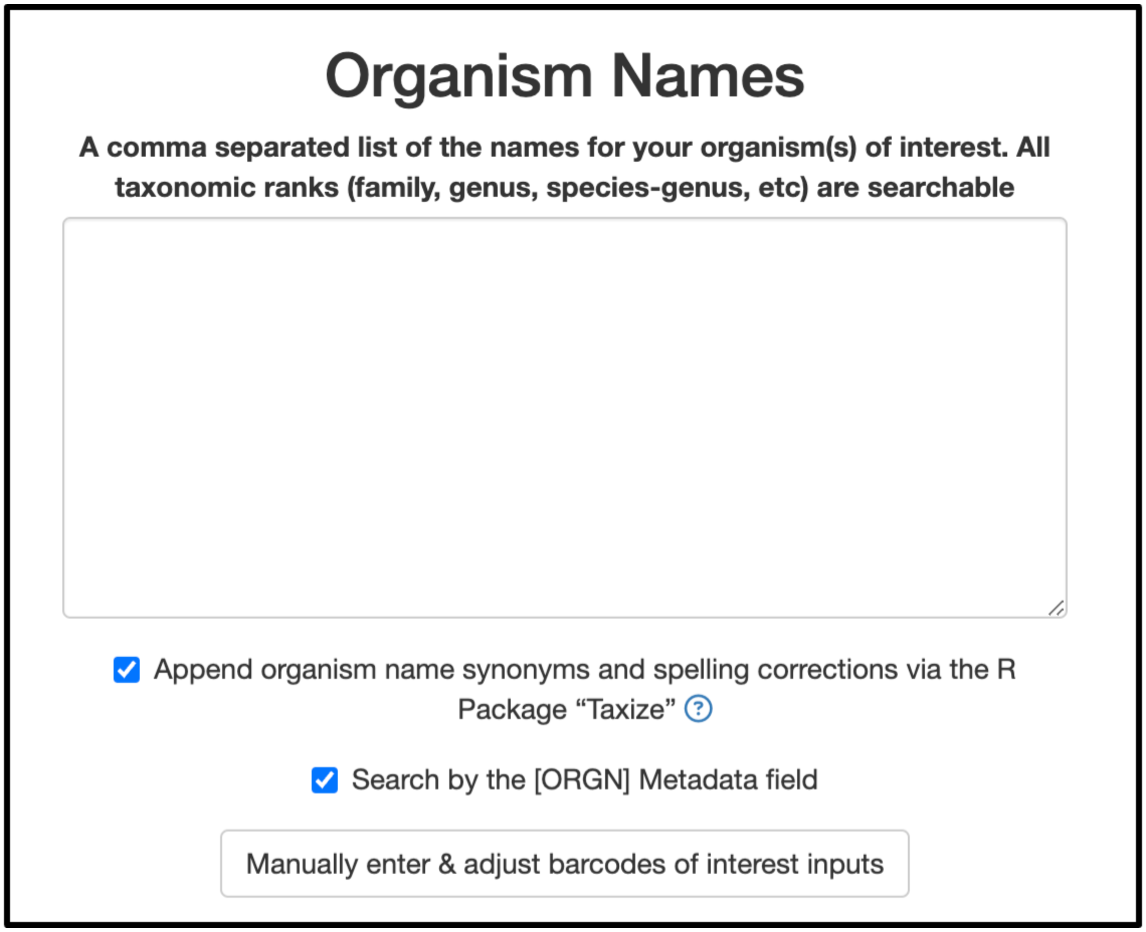

8.1 The Organism Names tab has 3 main objects to pay attention to (ordered from top to bottom):

1. **Big Text Box:** If a CSV was uploaded in the previous step, this text box will contain all the organisms contained in that file. Regardless, names can be manually added, removed, or modified by typing in the box. Organism names should be separated by commas, and leading or trailing whitespace will be ignored.
2. **Taxize:** If this checkbox is selected then spelling errors will be corrected, outdated organism names will be replaced, synonyms names for each organism will be added to the search. It is important to note that no user submissions are removed, meaning any changes will be appended to the search to prevent any data loss.
3. **[ORGN] Metadata:** If this checkbox is selected, the tool will only fetch entries where the Organism metadata field refers to one of the organisms in the search query. This means that entries that have missing or incomplete metadata will not be fetched. For instance, if an entry contains a reference to an organism of interest in its title but not in its organism metadata field, that entry will not be fetched by the tool. If the checkbox is not selected, organism names are searched for within all of the fields.

Once finished, click the "Manually enter & adjust Organism name inputs" button to proceed to the next step.

9 **Barcodes of Interest:** This tab in the NCBI pipeline is similar to the organism tab, but it is **instead** focused on barcodes.

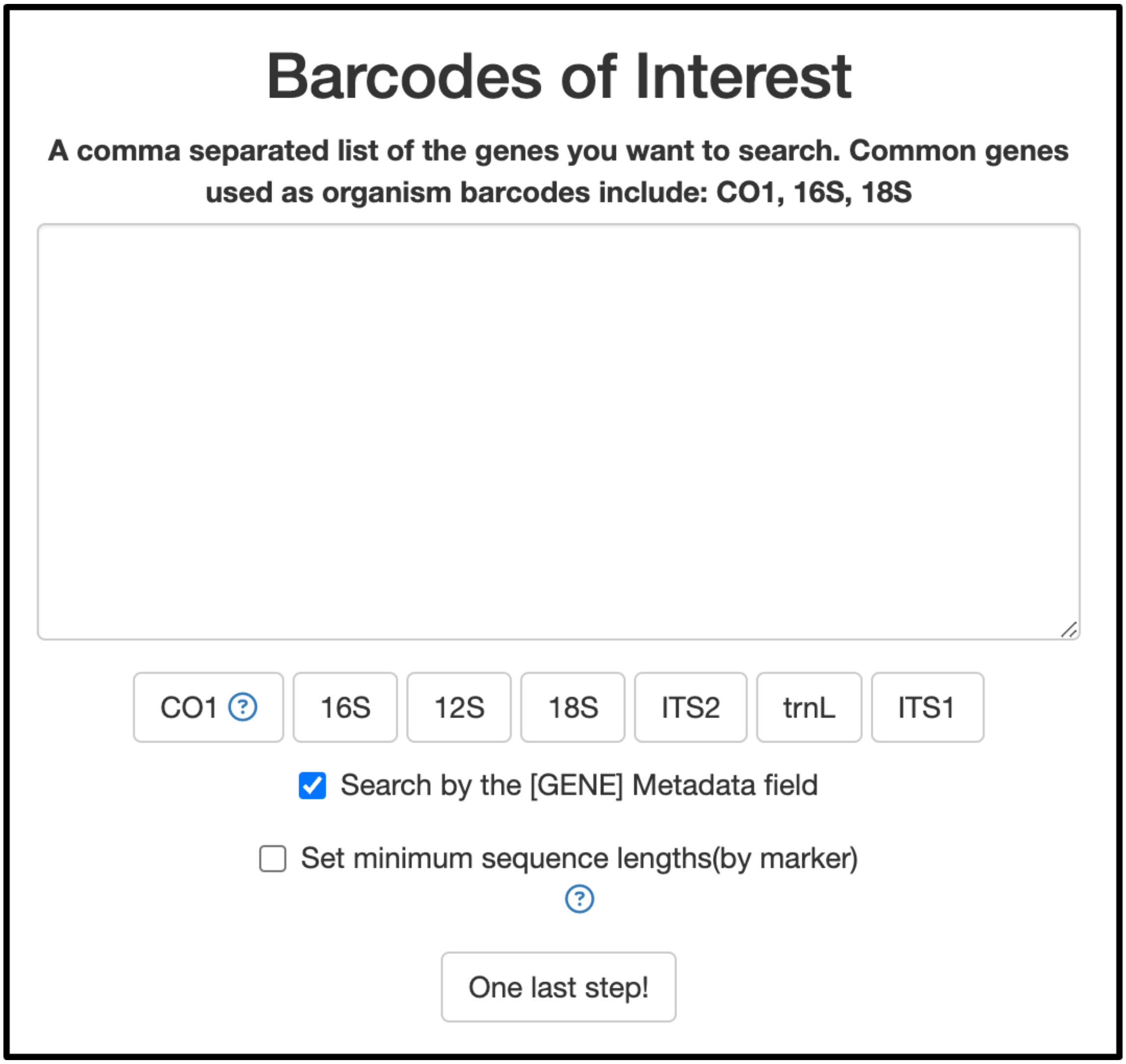


9.1 The Barcodes of Interest has 4 main objects to pay attention to (ordered from top to bottom):

- **Big Text Box:** If a CSV was uploaded in the previous step, this text box will contain all the barcodes contained in that file. Regardless, barcode names can be manually added, removed, or modified by typing in the box. Barcode names should be separated by commas, and leading or trailing whitespace will be ignored. Unlike the organism names screen, this screen gives the option of grouping together barcode names by using the following format:

```
( <Barcode name 1> + <Barcode name 2> + ...)
```

This feature will still search each barcode independently but display the merge the results together, which is useful for searching for barcodes that may have multiple synonyms. See the example below:

```
12S, CO1 + COI + COX1, 16S
```

- **Barcode buttons** (ex. CO1, 16S): These buttons are meant to quickly add barcodes to the textbox. The CO1 barcode in particular will also fill some commonly used synonyms (using the grouping feature mentioned above).
- **[GENE] Metadata:** If this checkbox is selected, the tool will only fetch entries where the gene metadata field refers to one of the barcodes in the search query. This means that entries that have missing or incomplete metadata will not be fetched. For instance, if an entry contains a reference to a barcode of interest in its title but not in its gene metadata field, then that entry will not be fetched by the tool. If the checkbox is not selected, barcode names are searched for within all of the fields.
- **Minimum sequence lengths:** If this checkbox is selected, a drop down menu will appear where minimum and maximum sequence lengths for each barcode can be individually specified. If it remains unselected, then no sequence length filtration occurs.

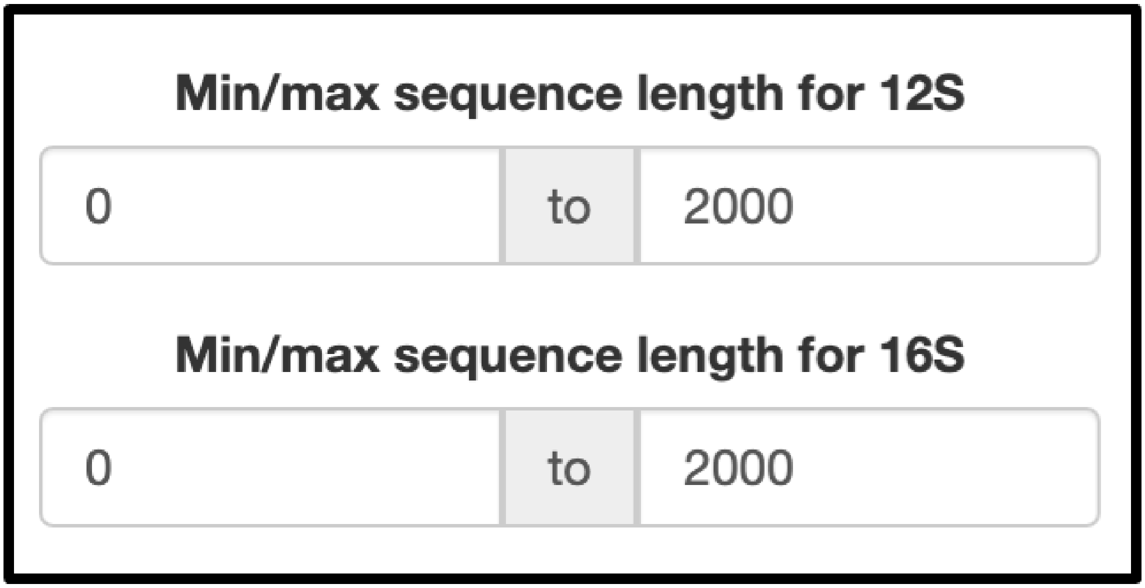


*Minimum sequence lengths example, these min/max boxes will appear for every barcode the user types into the textbox.*

Click the "One last step!" button to progress.

10 **Download Sequences Tab:** This tab in the NCBI pipeline is very simple. It contains a single text box which is used to specify the maximum number of sequences to download for each species/barcode combination in the search query. For example, to get 10 sequences of *Gallus gallus* for the barcode CO1, set this number to 10.

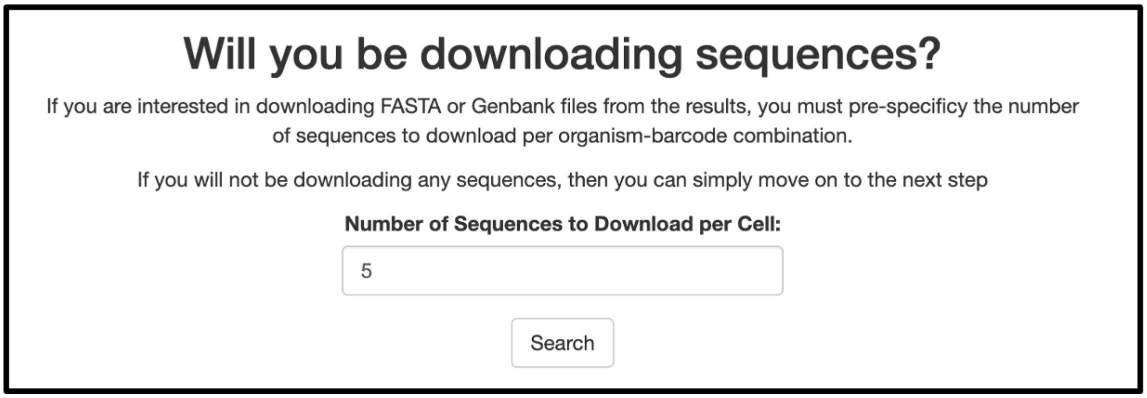


*Number of sequences to be downloaded for each species/barcode combination*

Finally, clicking the "Search" button will navigate to another page with a loading bar. Once the query finishes, a summary of the search results will be shown.

**Note**: that searches with over 50-100 species may take hours to complete. When accessing the tool online, it is important that the computer remains open and awake so that the session is not terminated by the server. When running the tool locally, it is fine to initiate a search right before going to sleep and leave the execution to run overnight.

11 **Summary Results Tab:** This tab will show a data table summarizing the search results. For each DNA barcode, the table contains the number of sequences found, the percent of total sequences found, the number of organisms with at least one sequence found, and the number of organisms with no sequences found.

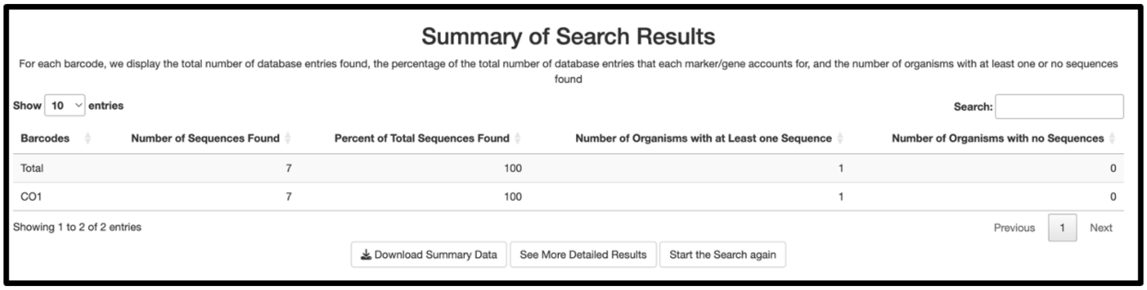


Summary data table together with the 3 main buttons this tab contains.

11.1 The summary results tab has 3 main buttons (from left to right):

- **Download Summary Data:** Click this button to download a CSV representation of this table.
- **See More Detailed Results:** Click this button to see a more detailed overview of the results.
- **Start the Search Again:** Click this button to start back at the beginning of the NCBI pipeline (**Step 7** in this protocol).

12 **Coverage Matrix Tab:** This tab contains two main tables, the top one is the coverage matrix table which contains the total number of entries found in NCBI with that species and barcode pair. The bottom table is the search terms table, which is included to assist in checking the

validity of the results by going to the NCBI website and pasting these queries into their online search tool.

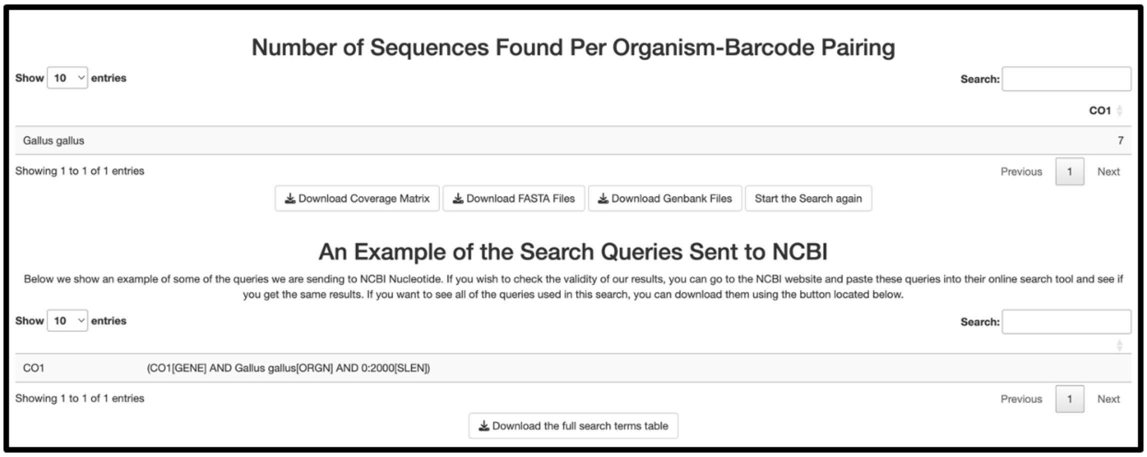


*Example of the Coverage Matrix Tab containing the coverage matrix table and the search queries table.*

12.1 **Coverage Matrix Table:** This table display the total number of entries/sequences found. Every row is an organism and every column is a barcode, the number that lies at every organism and barcode pair represents the number of entries found in the NCBI database.

Below the table are 4 buttons (left to right order):

1. **Download Coverage Matrix:** Click this button to download a CSV representation of this table.
2. **Download FASTA Files:** If this button is pressed, for every species and barcode pair the tool will download FASTA files for each of the entries found. The number of downloads of this button will depend on the number specified in **step 10**. For example, if there are 7 entries in NCBI and the number 3 was entered into the text box, the tool will download the first 3 entries out of the 7.
3. **Download Genbank Files:** If this button is pressed, for every species and barcode pair the tool will download GenBank files for each of the entries found. The number of downloads of this button will depend on the number specified in **step 10**. For example, if there are 7 entries in NCBI and the number 3 was entered into the text box, the tool will download the first 3 entries out of the 7.
4. **Start the Search again:** Click this button to start back at the beginning of the NCBI pipeline (**Step 7** in this protocol).

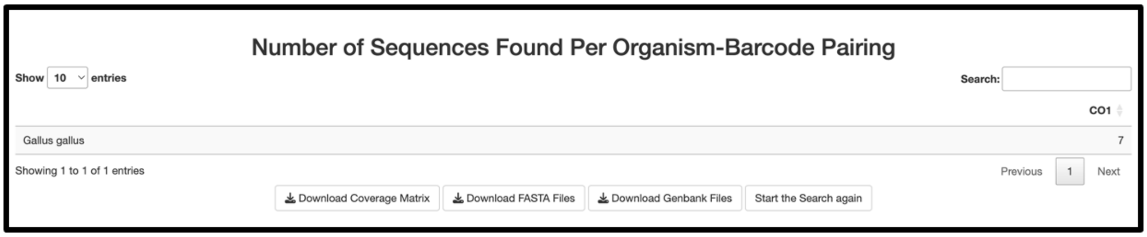


*Close up of the Coverage Matrix table together with all the buttons below it.*

12.2 **Coverage Matrix Table:** This table displays the exact NCBI database search terms that were used to query results for the first organism of interest in the search. This is included in the RSB as a validation tool to increase transparency and catch incorrectly generated search queries.

Click the "Download the full search terms table" button to download a CSV representation of this table.

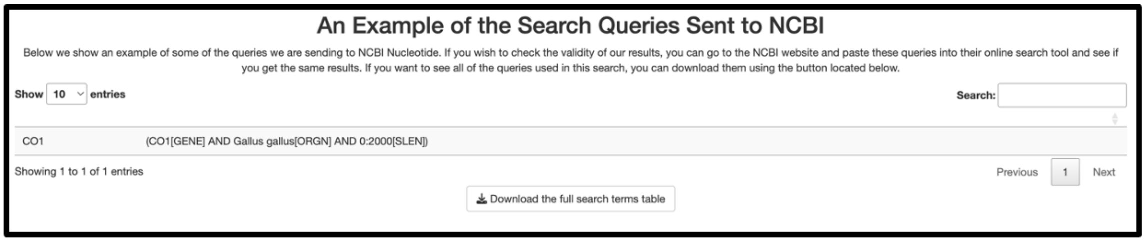


*Close up of the Search Terms table together with all the buttons below it.*

13 This is the the end of the NCBI pipeline. From here, either start a new search, import the FASTA files into your bioinformatic software/pipeline of choice, or continue to search other databases in this tool. We recommend looking into the BOLD pipeline, which supports de-duplicating NCBI entries.

### BOLD Tab

14 **Welcome Tab:** Using the navigation bar at the top, we can go to the BOLD section of the tool.

This tab has an identical structure to the NCBI welcome tab, the main difference is that CSV file for this pipeline need not have any barcodes, as it will only use the Organism columns of the CSV. If some of these buttons are unclear, check step 7.1 to see their functionality in the NCBI welcome tab.

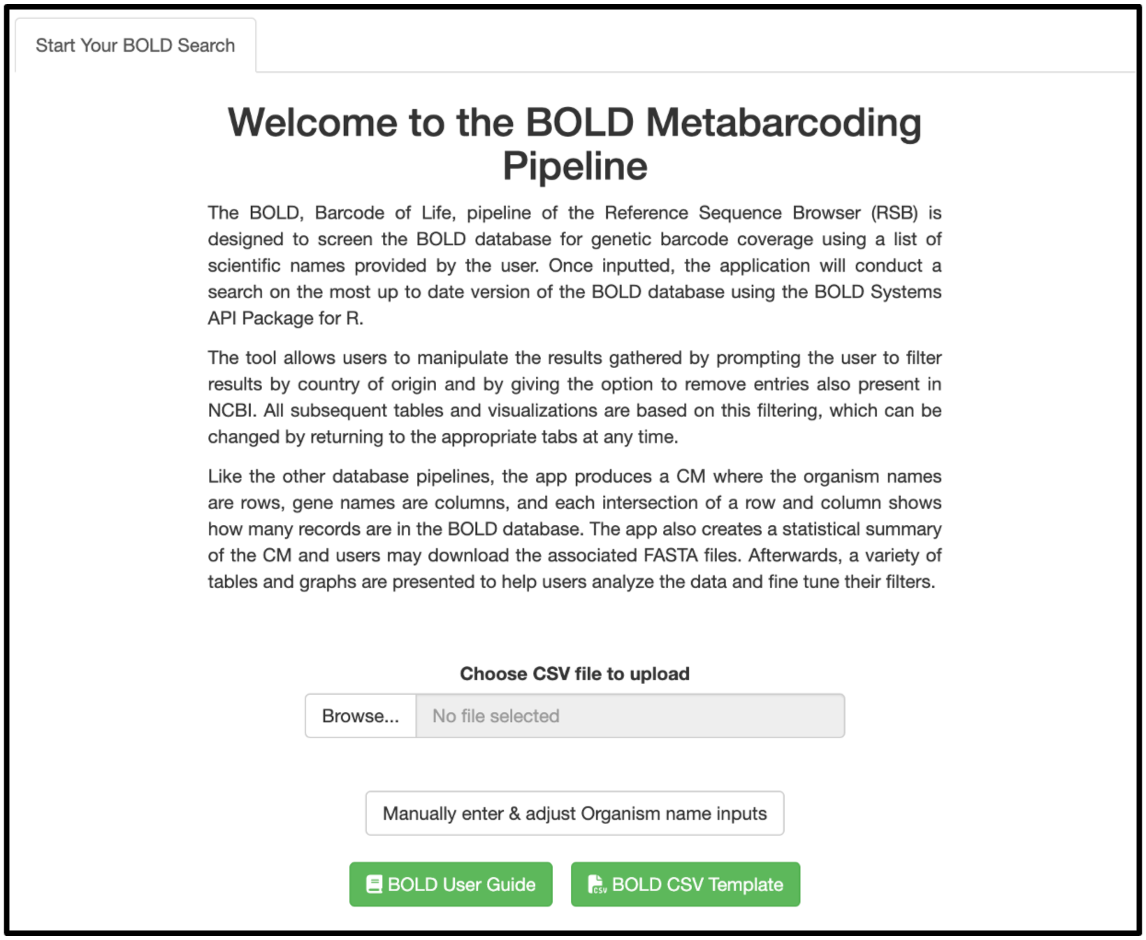


*Welcome page of the BOLD pipeline*

15 **Organism Names:** This tab is very similar to the NCBI tab. The only difference is that this screen lacks the [ORGN] Metadata checkbox because BOLD doesn't have the same metadata structure as NCBI Nucleotide. However, it still has the taxize setting, which works the same way as the one in the NCBI pipeline. Check step 6.1 to understand what the taxize checkbox does.

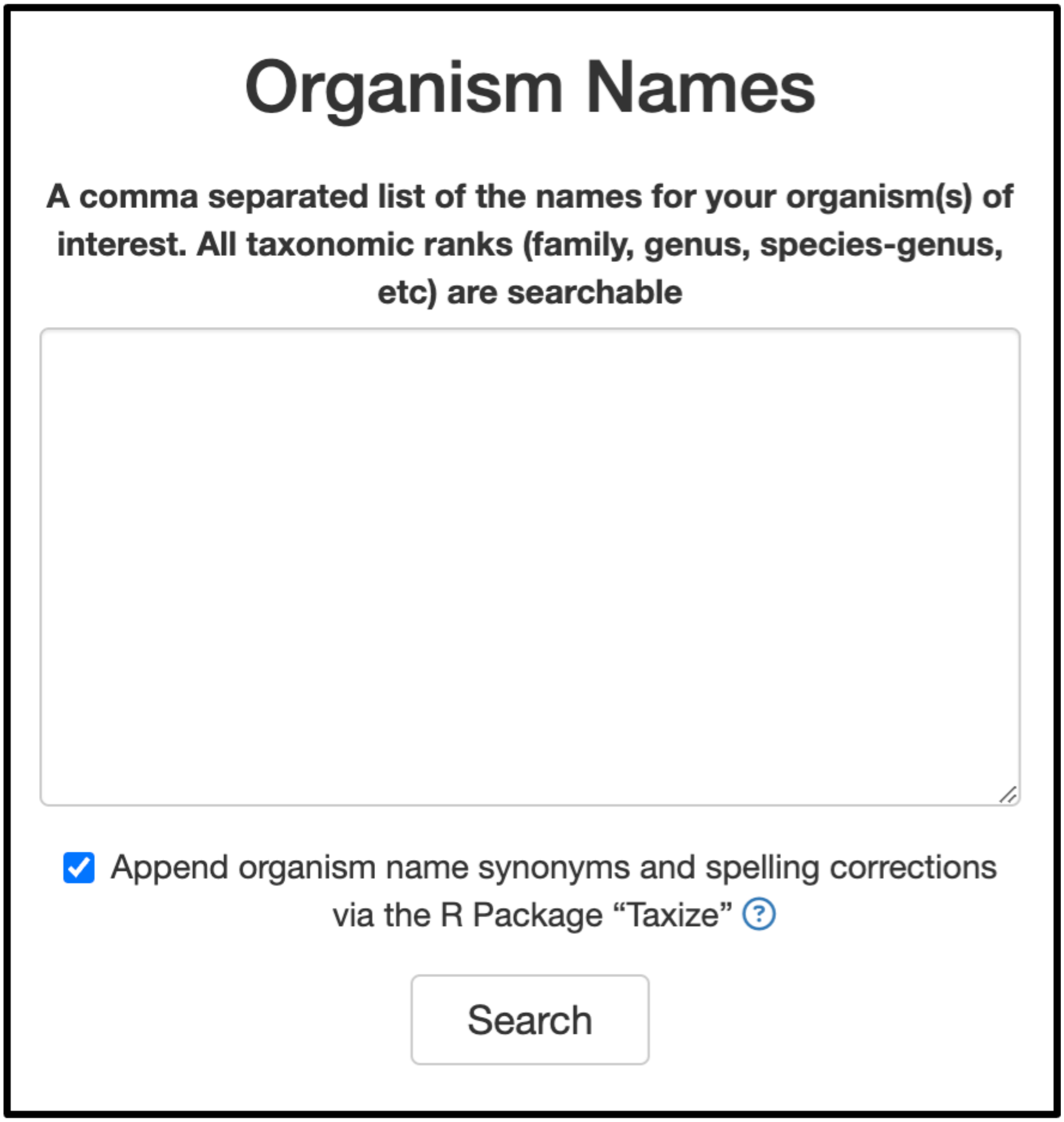


*Organism Tab of the BOLD pipeline*

Once all organisms of interest have been added, click the "Search" button.

16 **Missing/Invalid Metadata Tab:** The table in this tab lists organisms that were either not found in the BOLD database or yielded invalid results. If an organism does not appear in this table, it means there are results available in the BOLD database for it. Should the name of an organism in the table be appended with the (invalid) keyword, we recommend conducting a manual search to check if the issue stems from incorrect metadata returned by the BOLD database. Furthermore, please note that the results displayed in this table are unaffected by the filter settings.

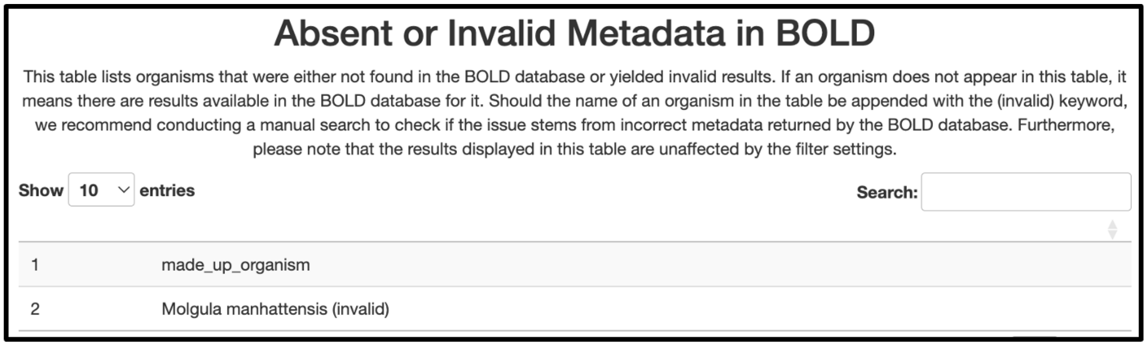


*Species not found in BOLD database table example*

17 **Filter Tab:** The BOLD filter tab is one of the most important in this pipeline, as it includes settings to filter by countries and a way to de-deduplicate NCBI entries.

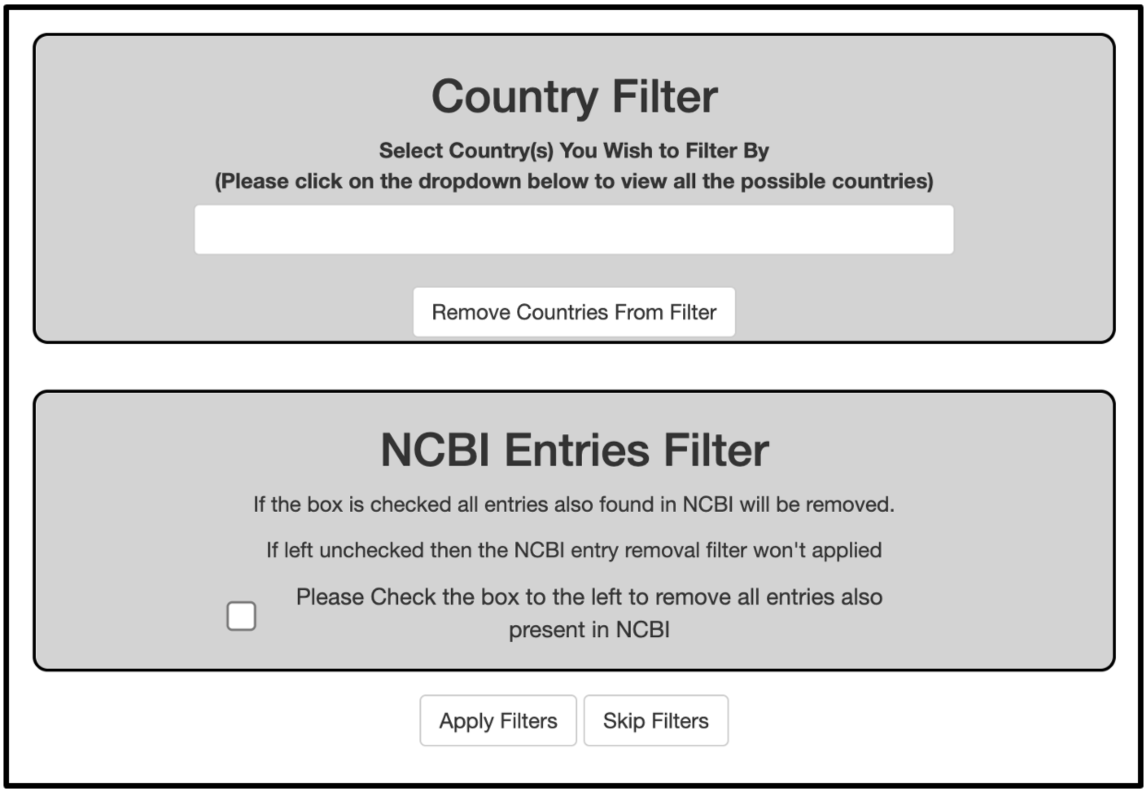


*Filter Tab example of the two filters countries and NCBI entries*

17.1 **Country Filter:** In order to use the country filter, click on the text box (marked in the image below with a red box). Doing this will open a dropdown menu with a list of countries found in the search. Choose any of the countries associated with geographic regions of interest. If filtering by geographic region is not necessary, simply leave it blank and click apply filter or skip filter.

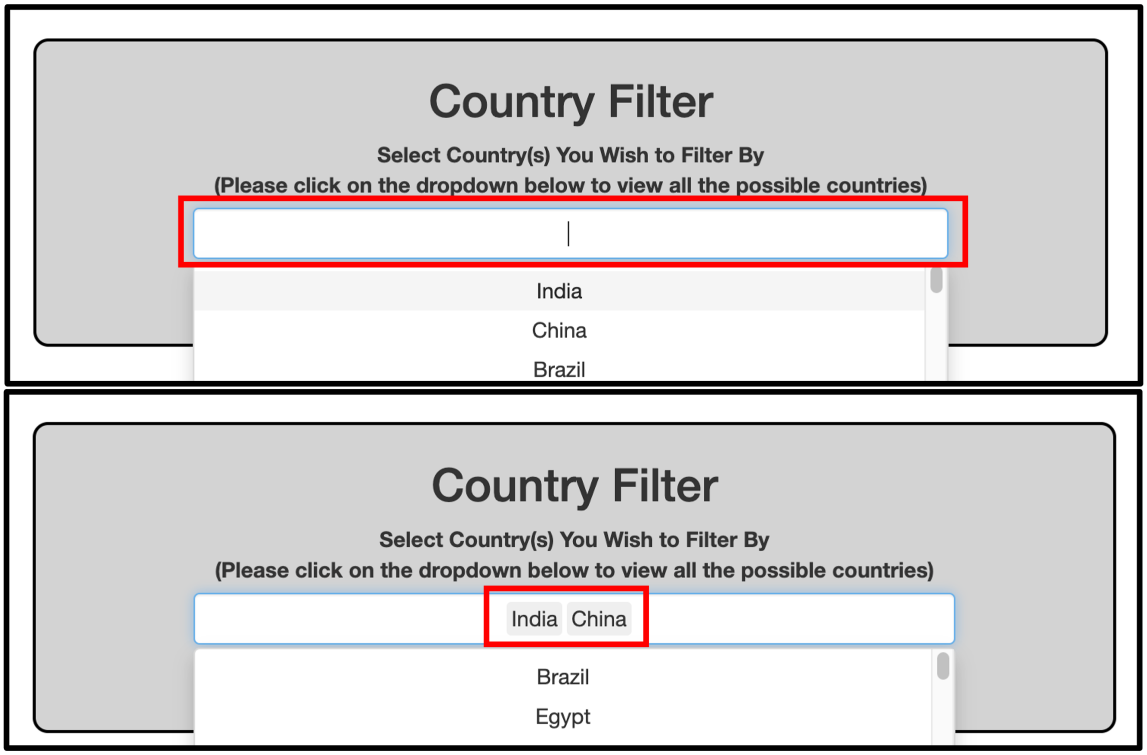


*Example of the dropdown menu and choosing countries to filter by in the BOLD Filter Tab*

17.2 **NCBI Deduplication:** If this checkbox is selected, it will remove any entries that are also found in NCBI. This allows you to only look at BOLD only entries instead of a mix of both databases as many NCBI entries are also found in BOLD.

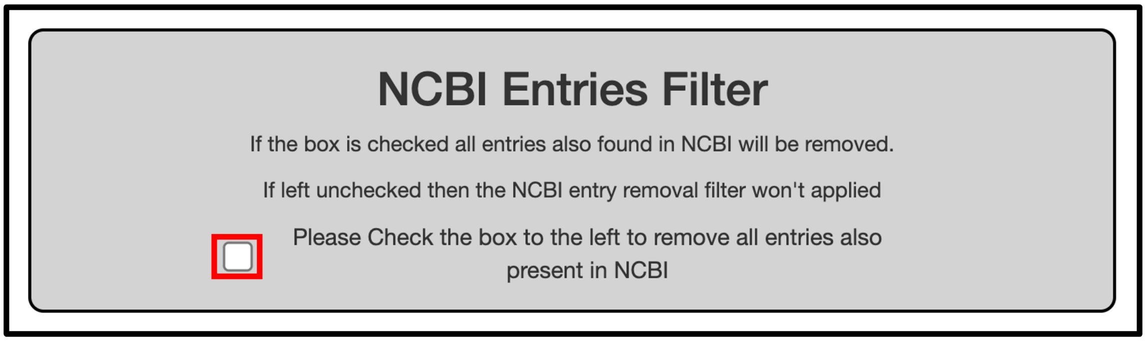


*NCBI deduplication checkbox*

18 **Results Section:** After clicking either the "Filters" or "Skip Filters" buttons, a lot of tabs will appear, each of them containing several ways to analyze the data and identify areas of missing information. Starting in this section, there will be no buttons to navigate from one tab to another. Explore them in whatever order makes sense in the given situation.

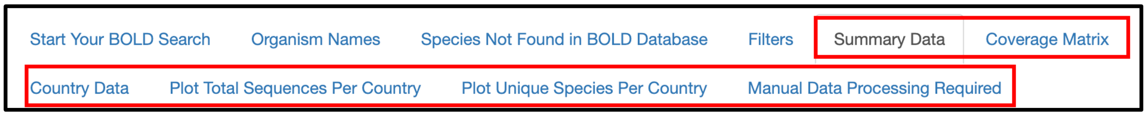


*Example of the all the tabs that show up once the BOLD filters are applied*

19 **Summary Data and Coverage Matrix:** These two tabs have exactly the same information as the ones in NCBI (covered in **steps 11 & 12**). The only difference is that the Coverage Matrix tab lacks a Genbank file download, since BOLD doesn't have those files.

20 **Country Data:** This tab has two main tables. Each of them are focused on the country information that was gathered in BOLD.

**Note**: As a reminder, BOLD doesn't ask the user to provide a set of barcodes to search; instead, it gathers all the barcodes that are available for the respective species in the BOLD database.

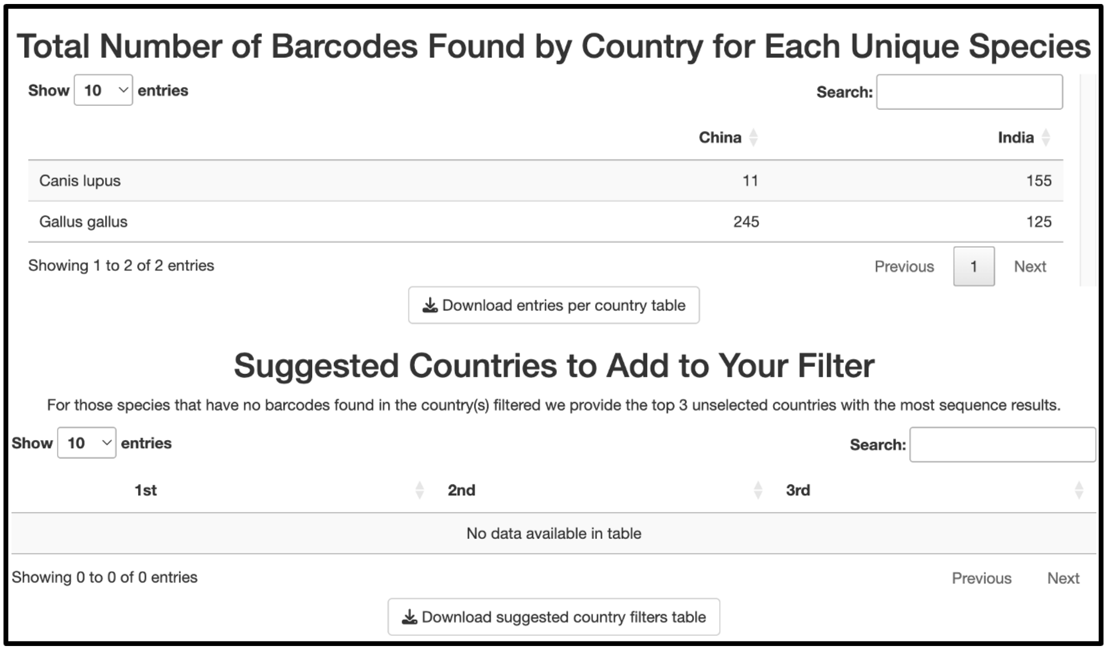

Total Number of Barcodes Found by Country for Each Unique Species

Show 10 entries Search:

| | China | India |
|---|---|---|
| Canis lupus | 11 | 155 |
| Gallus gallus | 245 | 125 |

Showing 1 to 2 of 2 entries Previous 1 Next

Download entries per country table

Suggested Countries to Add to Your Filter

For those species that have no barcodes found in the country(s) filtered we provide the top 3 unselected countries with the most sequence results.

Show 10 entries Search:

| 1st | 2nd | 3rd |
|---|---|---|
| No data available in table | | |

Showing 0 to 0 of 0 entries Previous Next

Download suggested country filters table

*Example of the two tables that are present in the country data table*

20.1 **Barcodes Found by Country:** The first table is not too dissimilar to the Coverage Matrix, as the numbers represent the number of sequences found in the BOLD database. However, instead of having barcode as the column it has countries. In other words, this table displays how many total barcodes were found in each country for each species.

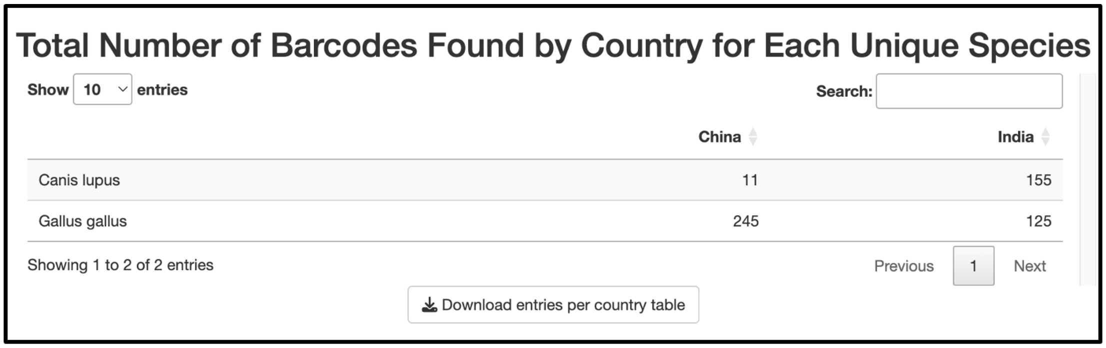

Total Number of Barcodes Found by Country for Each Unique Species

Show 10 entries Search:

| | China | India |
|---|---|---|
| Canis lupus | 11 | 155 |
| Gallus gallus | 245 | 125 |

Showing 1 to 2 of 2 entries Previous 1 Next

Download entries per country table

Example of the 1st table that appears in the Country Data Tab. Same as the one shown above but focusing only in 1 of the 2 tables that are created in this tab.

20.2 **Suggested Countries:** The second table only displays species that have no entries found under the applied filters and provides the top 3 countries with the most results. This table should help you add other countries to the filter if some of the species you are interested in have no results.

In **step 20.1**, we can see that the sequences for *Canis lupus* and *Gallus gallus* were gathered from India and China, the two countries we selected in the filter. Therefore, because there are reference sequences available for both organisms, the second table containing country suggestions will be empty.

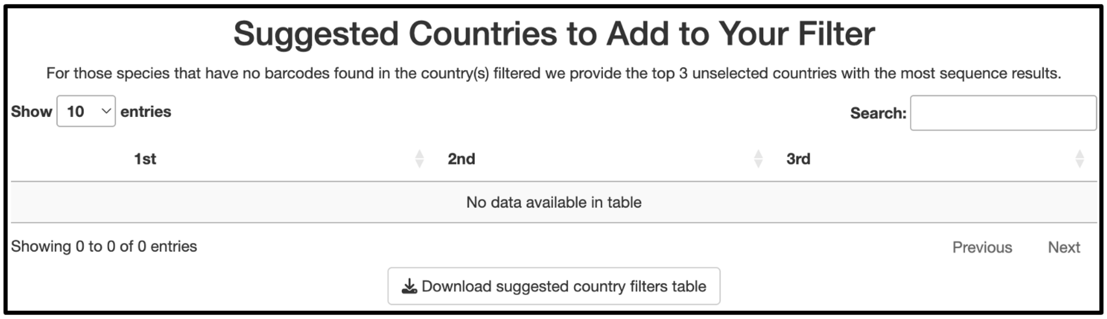

Suggested Countries to Add to Your Filter

For those species that have no barcodes found in the country(s) filtered we provide the top 3 unselected countries with the most sequence results.

Show 10 entries Search:

| 1st | 2nd | 3rd |
|---|---|---|
| No data available in table | | |

Showing 0 to 0 of 0 entries Previous Next

Download suggested country filters table

Example of the 2nd table that appears in the Country Data Tab displaying no results as both *Gallus Gallus* and *Canis Lupus* have results originating from China and India.

However, if you did not get any results for some species with the countries you filtered by, you would see three suggested countries that, from left to right, contain the most sequences for each species you are interested in.

### Suggested Countries to Add to Your Filter

For those species that have no barcodes found in the country(s) filtered we provide the top 3 unselected countries with the most sequence results.

Show 10 entries Search:

| | 1st | 2nd | 3rd |
|---|---|---|---|
| Canis lupus | No Country Listed | India | Australia |

Showing 1 to 1 of 1 entries Previous 1 Next

Download suggested country filters table

Example of the 2nd table that appears in the Country Data Tab displaying results as *Canis Lupus* had no reference sequences from the countries filtered for the purposes of the example.

21 **Plots:** The next two tabs present two plots to help further analyze the results of the BOLD database.

21.1 **Tree Map Sequences per Country**: The first plot is a tree map that shows the distribution of the total sequences found per country, and is meant to display how many sequences are in each country.

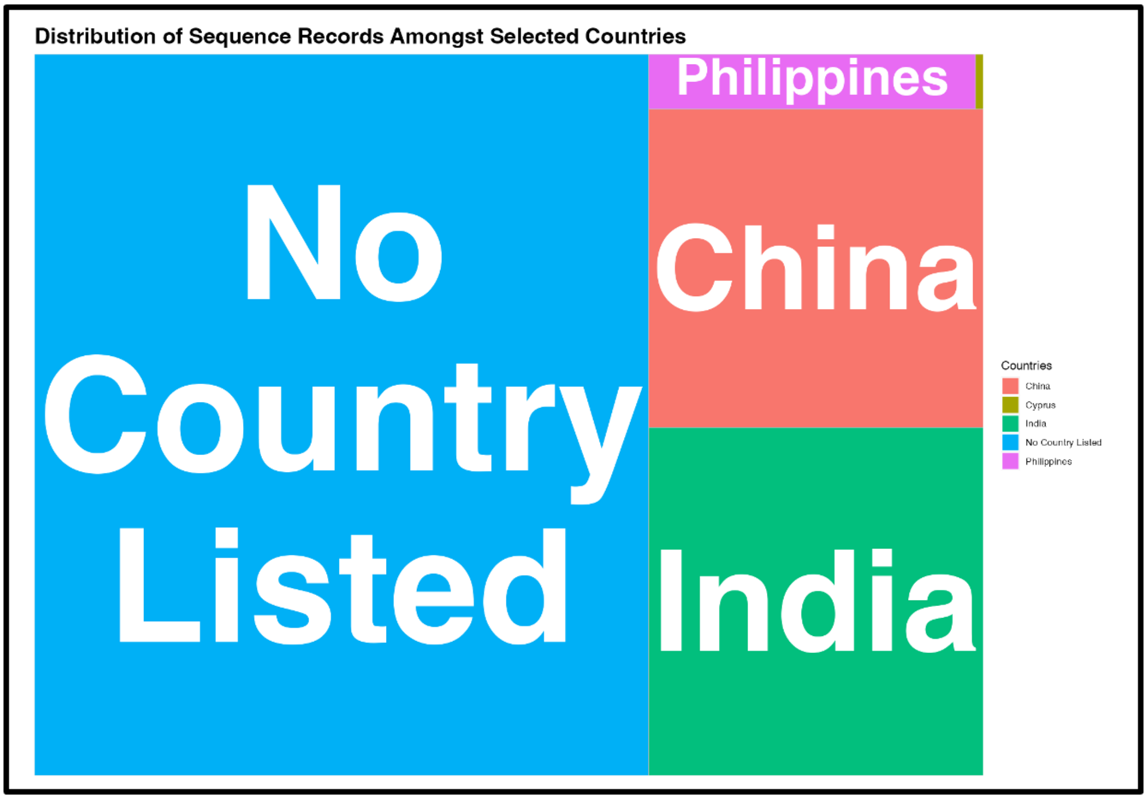


*Tree map example of the BOLD plots displaying graphically how many sequences each country has*

21.2 **Bar Plot Unique Species per Country:** While it is useful to see how many total sequences each country has, it is also important to see how many unique species each country has reference sequences for. Ultimately, the results may be distributed such that a single country that has 90% of the total sequences found but only has reference sequences for 1 of the many organisms you are interested in.

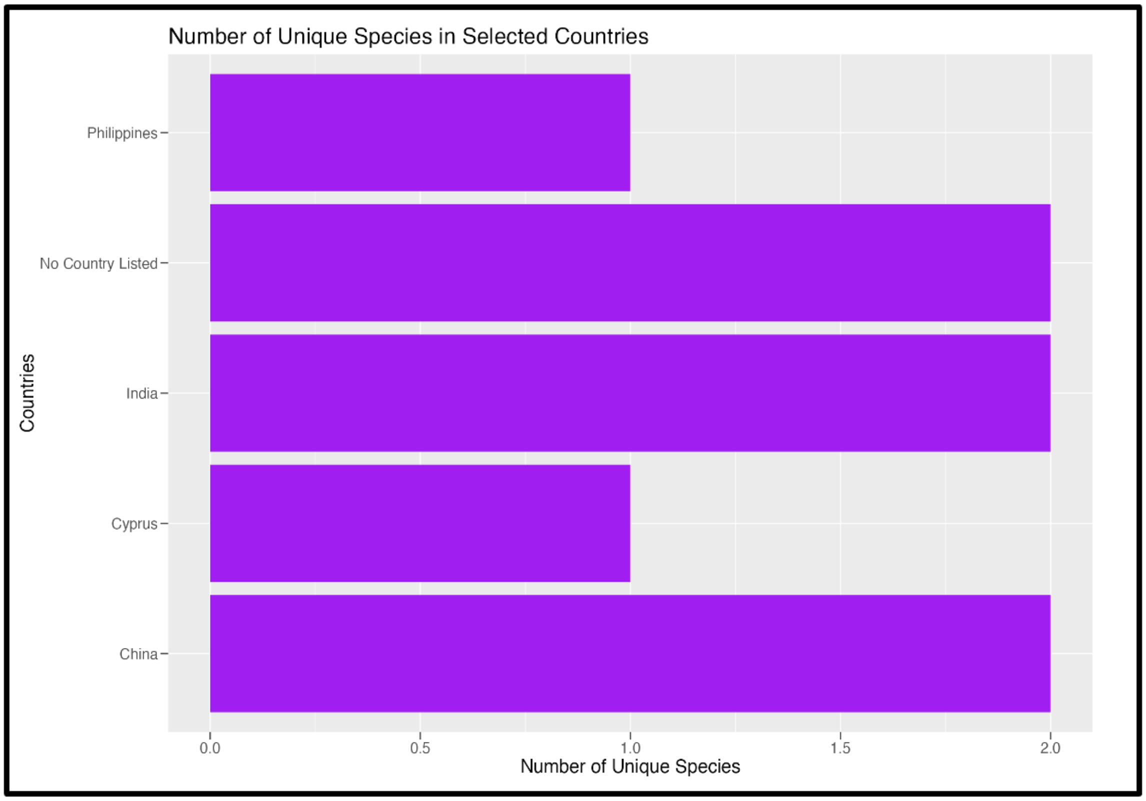


*Bar plot example of the BOLD plots displaying graphically how many unique species each country has*

22 **Manual Data Processing Required:** Not all entries in BOLD have sequence attached to them. For the sake of transparency, this table contains a listing of all such entries for each species in the search. It may be worth manually examining these entries in BOLD to see what information they contain.

| | Entries where Barcode was NA |
|---|---|
| Bythotrephes longimanus | BYTHO114-22, CAISN938-13, NJCGS015-09, NJCGS016-09 |
| Canis lupus | NOMAM074-09, ZSILG432-22, NOMAM145-17 |
| Caprella mutica | BNSB040-21, DUTCH667-19, DUTCH668-19, GBCMA12312-16, QHAK022-20, QHAK026-20, QHAK032-20, QHAK034-20, QHAK037-20, SWEMA466-15, SWEMA864-15, BNSB038-21, BNSB039-21, BNSB035-21, BNSB036-21, BNSB037-21 |
| Cordylophora caspia | ANNMO567-20, SWEMA984-15 |
| Crepidula fornicata | BNAGB664-14, LITOR363-10, LITOR364-10, LITOR365-10, SWEMA033-15 |

*Example of the BOLD manual data processing required*

### CRUX Tab

23 **Welcome Tab:** Using the navigation bar at the top, we can go to the CRUX section of the tool.

This tab has an identical structure to the NCBI welcome tab, the main difference is that CSV file for this pipeline need not have any barcodes, as it will only use the Organism columns of the CSV. If some of these buttons are unclear, check step 7.1 to see their functionality in the NCBI welcome tab.

**Note:**

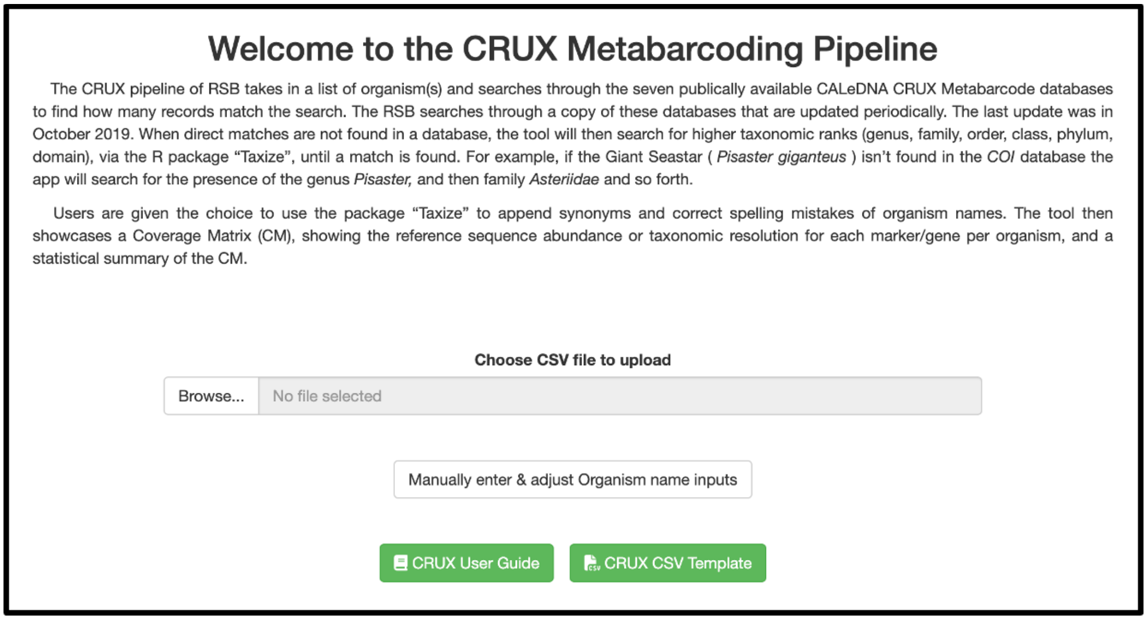


*Welcome section of the CRUX tab*

24 **Organism Names:** Clicking the "Manually enter & adjust Organism name inputs" button it will navigate to the CRUX Organism Names tab.

This tab is very similar to the other Organism Names tab as it still has the text input box and the taxize checkbox. However, there is one more checkbox option one can select which checks for any homonyms found in your dataset.

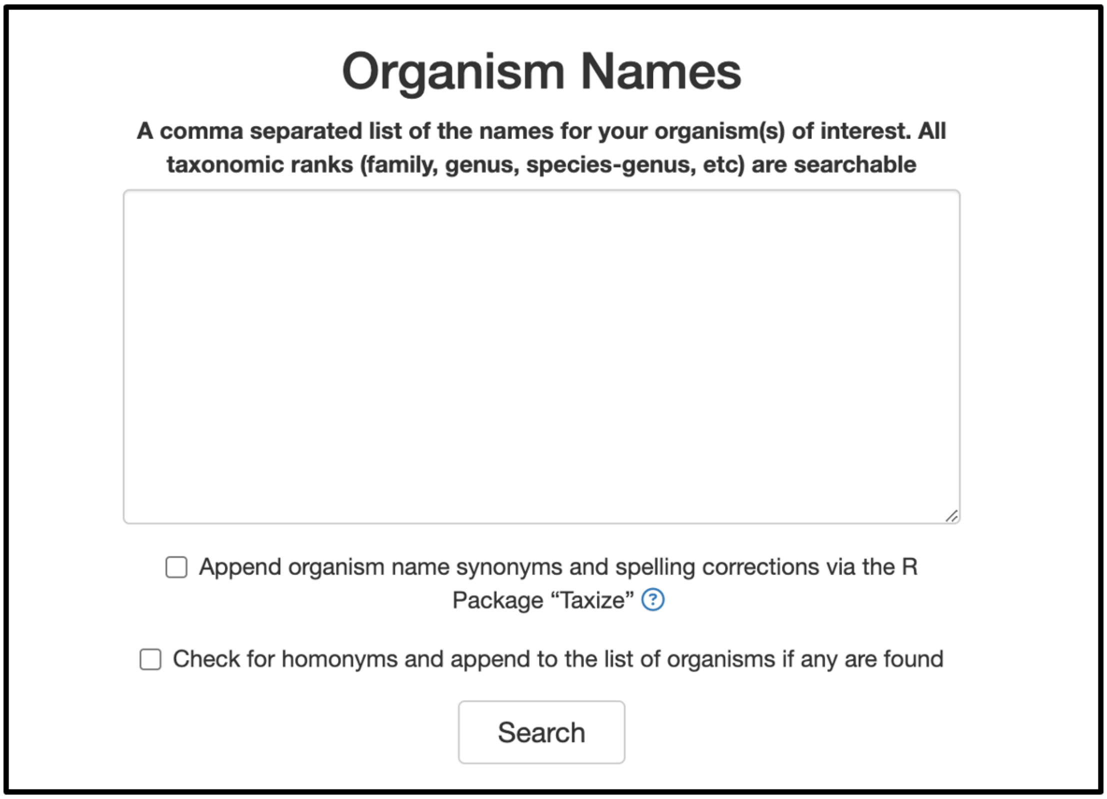


*Organism Tab of the CRUX pipeline*

24.1 **Homonym Example:** If the homonyms checkbox is selected, and you were to input an organism that has a homonym (i.e. a taxon that is identical in spelling to the name of a different taxon). In the example below, the taxon *Asterina gibbosa* is the name of both a cushion starfish and a fungus.

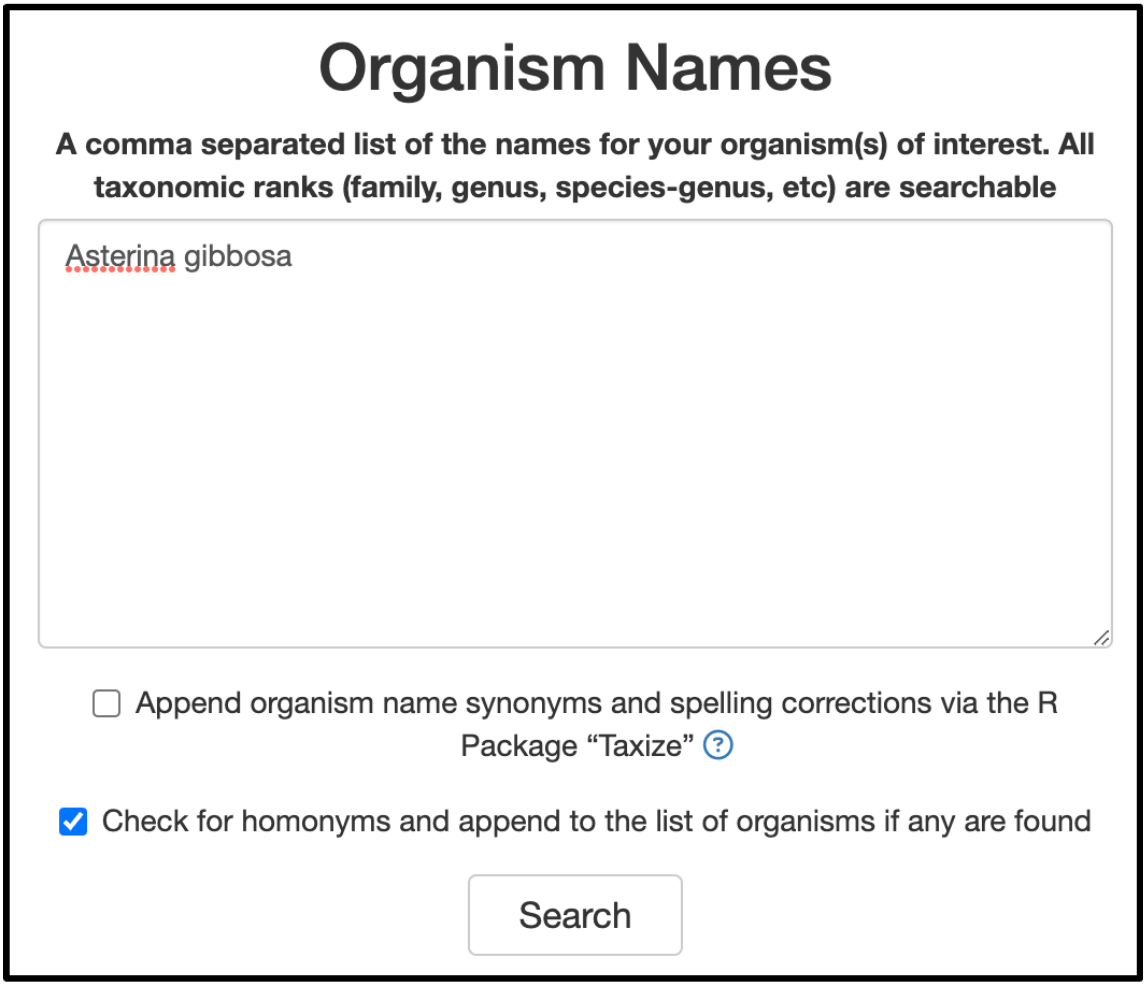


*Example of a species genus that has homonyms and the homonym checkbox selected*

After clicking search, notice that the search outputs 3 different rows in the Coverage Matrix table. The first entry is the one the user wrote down(*Asterina gibbosa*). The second and third entries are the two types of *Asterina gibbosa*: the starfish and the fungus.

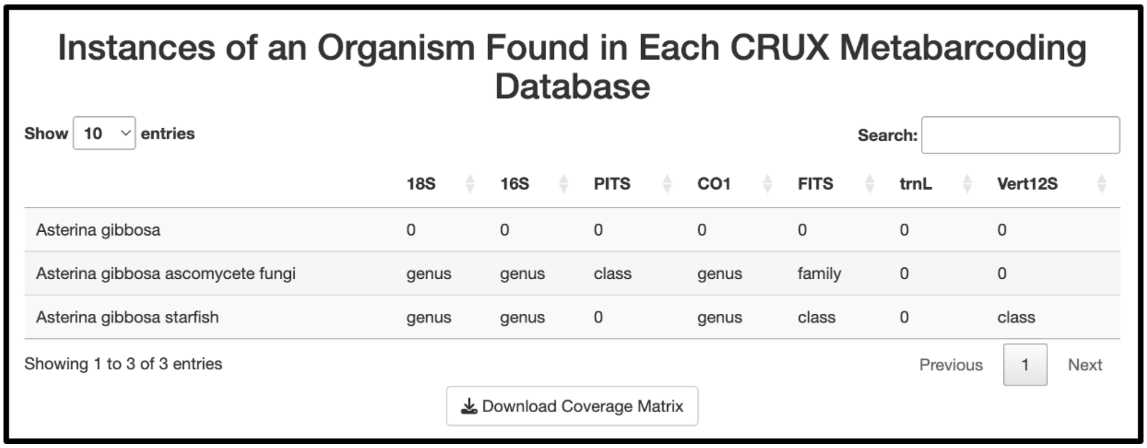

Instances of an Organism Found in Each CRUX Metabarcoding Database

Show 10 entries Search:

| | 18S | 16S | PITS | CO1 | FITS | trnL | Vert12S |
|---|---|---|---|---|---|---|---|
| Asterina gibbosa | 0 | 0 | 0 | 0 | 0 | 0 | 0 |
| Asterina gibbosa ascomycete fungi | genus | genus | class | genus | family | 0 | 0 |
| Asterina gibbosa starfish | genus | genus | 0 | genus | class | 0 | class |

Showing 1 to 3 of 3 entries Previous 1 Next

Download Coverage Matrix

*Example of the results of searching a species genus that has homonyms and having homonym checkbox selected*

25 **Summary data:** This tab has exactly the same information as the one in NCBI that was covered in **step 9**.

26 **Coverage Matrix:** This table in CRUX database pipeline is slightly different from the tables in **steps 12 & 19**. The rows still represent an organism, but the columns represent one of the public CRUX databases, and values in the table may not always be numbers. When the tool does not find direct matches in a database, it will instead search for higher taxonomic ranks of that organism until it finds a match. If the RSB performs lower-resolution searches, it will output the rank at which it found a sequence instead of displaying a numerical value. The tool works its way up the following taxonomic ranks: species, genus, family, order, class, phylum, and domain.

#### Instances of an Organism Found in Each CRUX Metabarcoding Database

Show 10 entries Search:

| | 18S | 16S | PITS | CO1 | FITS | trnL | Vert12S |
|---|---|---|---|---|---|---|---|
| Canis Lupus | genus | genus | phylum | genus | order | 0 | genus |
| Gallus gallus | 13 | 193 | phylum | 233 | class | 0 | 186 |

Showing 1 to 2 of 2 entries Previous 1 Next

Download Coverage Matrix

Example of the CRUX Coverage Matrix, as described above some results display a rank instead of a number as they didn't have direct results in the database and had to go up.

### Questions & Issues

27 **Questions:** For any further questions about the tool, see our paper (link can be found in the Github) or contact us directly. If you are using this tool for a study, we would love to hear feedback on future features that could be added to it.

**Issues:** If any issues or bugs come up while using this tool, please open an issue in the GitHub and let us know. The link to our Github is: https://github.com/SamuelLRapp/BlueWaltzBio.
